# Daily activity and visual discrimination reflects the eye organization of weaver ant *Oecophylla smaragdina* (Insecta: Hymenoptera: Formicidae)

**DOI:** 10.1101/193243

**Authors:** Monalisa Mishra, Snigdha Bhadani

## Abstract

The currently studied ant *Oecophylla smaragdina* is arboreal in nature. It shows unique property in terms of its nest building behavior using leaves of the tree. The ant uses its eye to locate the food and nest. Eye types vary among ants living in the different habitat. In this context, the structure of the eye, daily activity and the foraging behaviour of *O. smaragdina* are missing from the literature. For the first time, the current study discovers: (1) the ant *O. smaragdina* forage in fully lighted condition. (2)The eye structure shows that the eye is adapted to a diurnal lifestyle.(3) The daily activity is proportional to the temperature of the surroundings. The study significantly correlates the role of vision in the foraging behaviour of *O. smaragdina*. The daily activity is further associated with the surrounding light and ambient temperature. The current study uncovers the structure of the eye and eye-related behaviour of the animal not described in earlier studies.

## Introduction

Ants are the most dominating and successful insects found in various niches (Culver, 1972; Davidson, 1977; Mezger and Pfeiffer, 2010; Torres, 1984). The compound eye is one the parameter which makes them successful to live in various niches (Menke et al., 2009; Moser et al., 2004; Narendra et al., 2013). The compound eye organization varies within species, niches and with activity time (Moser et al., 2004; Narendra et al., 2013). Thus, ant’s eye is a nice model to study the functional physiology of the animal. Ant’s eye irrespective of various activity time and niche studied by various authors (Culver, 1972; Torres, 1984). The ant’s active during daytime possess different structural arrangement than the one active during night time. In general, the diurnal ants possess smaller facet diameter and rhabdom size than the one active during night time (Moser et al., 2004). For example, the diurnal ant like *Melophous bagoti* possesses the facet (lens of the eye) diameter 19μm (Schwarz et al., 2011). In nocturnal ant *Myrmecia pyriformis,* the facet diameter is twice than the size of *M. bagoti* (Greiner et al., 2007). Besides facet, the light sensing organelle or the rhabdom also increases its size approximately four times than nocturnal ant *M. pyriformis*. Adaptations in night active ants are meant to capture every stray of light in a photon-starved condition. Compound eye further achieves the resolution of the eye by altering the number and size of the facets with it’s activity time. Thus the size of the eye depends more on the activity time rather than the body size of the animal (Greiner et al., 2007; Moser et al., 2004).

The ant *Oecophylla smaragdina* is famous for its arboreal nature and its nest building property using the leaf (Fiedler and Maschwitz, 1989; Holldobler, 1983; Schlüns et al., 2009). The worker ants are known to forage in a group and carry egg, larvae and young from one tree to another (Devarajan, 2016). The eye has a very crucial role in determining the location of the nest and behavior of the animal. In this context, the eye ultrastructure, activity rhythm and foraging behavior of this ant are missing from the literature. The current study deciphers the eye of the animal with its visual activity not described in earlier studies.

## Materials and Methods

### Activity Rhythm

To analyze the activity rhythm of *O. smaragdina*, a nest on neem and jackfruit tree was chosen which is located at National Institute of Rourkela, Rourkela campus, Odisha, India (latitude and longitude of 22.25N and 84.90E respectively). Their rhythmic activity was visually monitored on a 15 hours daily basis i.e. from 4:00 am in the dawn till 8:00 pm dark. For proper visualization of ants activity, an imaginary circle of diameter approximately 50 cm was assumed around the nest entrance. The numbers of inbound and outbound activity of these ants were calculated at an interval of two minutes for every two minutes. This activity was monitored for 30 days. During this experiment, the environmental temperature was also correlated with the activity of *O. smaragdina.* The graphs were plotted to know the number of inbound/outbound ants at a particular time of the day.

### Allometry and Eye Structure

#### Sample collection

To view the organization of ant’s eye, ants were collected near the nest in a plastic bottle having holes and carried to the laboratory.

### Measurement of thorax length

To measure the thorax length, ants were anesthetized by putting cotton, soaked with diethyl ether, inside the bottle. Ants were imaged along with a scale under a stereo microscope. Images were transferred to Image J and thorax length was calculated.

### Scanning electron microscopy

Ant heads were cut using a razor blade and fixed with 4% paraformaldehyde at 4°C for overnight. Next day, heads were washed with phosphate buffered saline (PBS) and dehydrated using serial dehydration of ethanol to remove the moisture content. The dehydrated heads were kept in a desiccator for overnight. Next day, heads were mounted on a stub having double adhesive carbon tape on it with the help of a stereo microscope. Heads were coated with platinum for 3 minutes with the help of a sputter coater (Q150RES, quorum Tech). Coated heads were imaged under FESEM (Field Emission Scanning Electron Microscope (Nova Nano SEM 450, FEI) operated voltage of 10 kV. All the images were transferred to Image J to calculate the various parameters of the eye.

### Measurement of surface area of eye

SEM images of the whole eye were used to measure various parameters. The length (*Cl*), width (*Cw*) of the compound eye and number of ommatidia were calculated. The total surface area of the eye was calculated putting the values of *Cw* and *Cl* in the equation mentioned below, where A is the area of ellipse

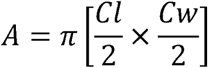

### Calculation of mean ommatidial diameter

This parameter gives the description of cornea sampling density and the light sensitivity of eye respectively.

### Calculation of interommatidial angle

The frontal view image of ant’s head was used for this analysis. Three points were drawn on the external perimeter of the eye using Image J. A circle was made passing through these points in such a way that the boundary of the circle touches the boundary of the eye. One horizontal and vertical line was drawn in such a manner that they intersect at the center and make an angle of 90°. For calculating the interommatidial angle (φ), the desired number of ommatidia was calculated from the eye’s perimeter (Meyer-rochow and Mishra 2007). For the current study, six numbers of ommatidia were calculated, and two lines were drawn from the outer boundary of the eye to the center containing the counted ommatidia using Image J software. The angle subtended by the lines at the center was calculated and the *mean ommatidial angle* was calculated by dividing the measured angle with a number of ommatidia calculated. The above steps were repeated for different portions of the compound eye to eliminate the error.

### Ommatidial diameter

The *ommatidial diameter*, D was obtained by drawing a line crossing a row of 10 ommatidia in the vertical or horizontal plane using Image J software and dividing the length by 10. The *mean ommatidial diameter*, D, was calculated for 3 times from different portions of the eye by choosing the areas randomly.

### Eye parameter

The eye parameter, *p* gives the relationship between the sensitivity and resolution of the eye. It was calculated as *p = D.* Δφ.

### Visual discrimination

The ants were taken to the laboratory to a constant environmental temperature of 32° C. Ants were kept in dark for six hours. Two choice assays were performed on them in an experimental device designed in wood. The wooden device was made to give familiar habitat to the ants. The device comprises of 3 wooden planks having a hole of diameter 2.5 cm, vertically mounted on a wooden base. Out of the three planks, two of them i.e. the sides ones are fixed (20x7 cm) on base and the middle one is movable (24.6x7 cm) and can be taken out. Two plastic tubes of size (2.3x9.3 cm) diameter were fitted in the holes, with an angle of 180°. The dark-adapted ants were kept inside central part of plank and allowed to choose various stimuli in either arm of the tube within the device. A sum total of 50 ants were used for the visual discrimination analysis.

#### Visual Stimuli

Sum total of three different visual stimuli was given to the ants. Stimuli were given to the ants by giving choice in different tubes of the device (1) light/dark (2) small field (upward and downward isosceles triangle) (3) large field (four black and five white stripes were drawn horizontally and vertically). At the end of each stimulus, the number of ants in each tube was analyzed.

For light/dark assay, dark-adapted ants were transferred to the experimental setup in a dark room. A beam of light was flashed from the base of one tube while the other tube was kept dark. Ants were further allowed to choose between two different (small-field) visual stimuli. Two small isosceles triangles of side 1cm each were made. One triangle was put pointing upward and the other pointing downward direction to different tubes of the above-mentioned device. Ant’s choice was calculated and plotted in the form of a graph. For the large field stimuli, 4 black stripes and 5 white stripes were drawn alternatively. In one pattern the lines were drawn horizontally and the other vertical lines were drawn on a white sheet of paper. The stripes are in 4mm width and a single stripe subtends a visual angle of 2.46°. The small visual stimuli offered a much smaller area as compared to large stimuli. This experiment was repeated for six times. Based on the result the mean value was plotted in the form of a graph.

## Results

### Activity rhythm

The activity rhythms of the *O. smaragdina* were visually monitored in natural conditions and the number of inbound and outbound ants was calculated. The inbound and outbound activity was further correlated with ambient temperature (Fig. 1). According to the record, the activity starts the early morning at 4:50 am when the temperature was around 32° C. During this time, all the ants returned to their nest. Around 5:00 am the ants started coming out of their nest. The inbound and outbound activity increases progressively with time and attains a peak around 7:00 am (Fig. 1). The temperature recorded was around 39° C. With the increase of temperature the activity increases up to 40.5° C. The activity decreases and stops completely at 9:30 am when the surrounding temperature increases up to 40.5° C. No activity is seen after that time since the ambient temperature was very high. Ants prefer to stay in the nest during that time. As the temperature decreases up to 39° C, their activity started around 5:30 pm. It was observed that their outbound activity is more than their inbound activity. The activity stopped around 6:25 pm. These observations establish a relation that the rhythm activity of *O. smaragdina* ants is dependent on temperature. Since no activity seen when there is less light, probably the eye is adapted towards the light condition.

**Figure 1.**
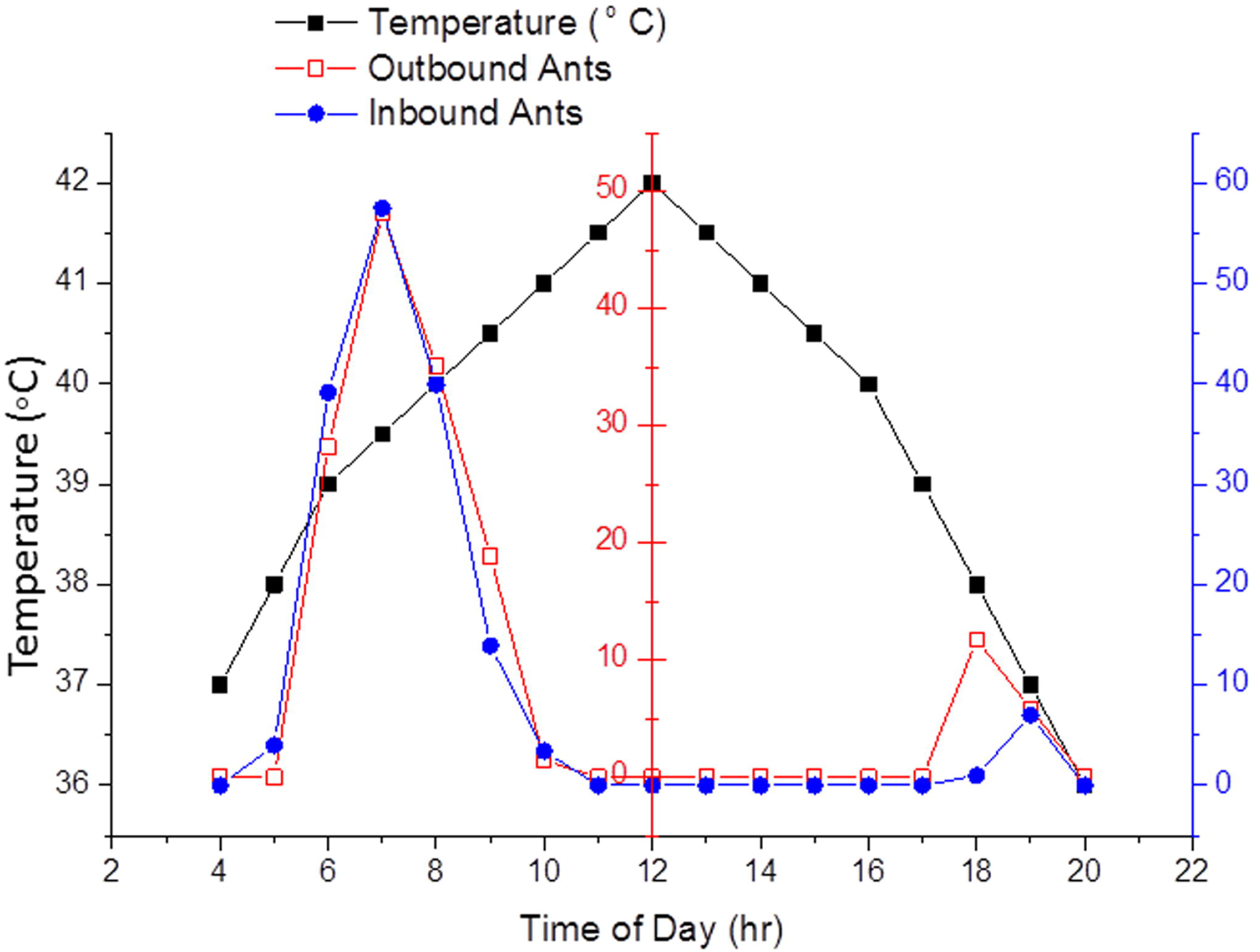
Activity rhythms of an ant*O.smaragdina* as a function of climatic temperature. Variation of inbound and outbound activity with temperature is depicted in this figure.

### Allometry and eye structure

The morphological characteristics of the compound eye were analyzed from FESEM images. The eye is spherical in shape in lateral view (Fig. 2A). Both the eyes are well-separated from each other (Fig. 2B) and the distance between the two eyes is 955.506 ±5 μm. The diameter of the eye is estimated to be 531.612 ± 5 μm. Each eye has approximately 700-800 facets or ommatidia. The shape of the eyes varies depending on the angle through which it is observed (Fig. 2C, D). The ommatidia which are present in the central part of the eye are hexagonal in shape (Fig. 2E). The peripheral ommatidia are an irregular hexagon, pentagon or square in shape (Fig. 2F). The interommatidial hair is randomly distributed throughout the compound eye and found to be more at the central part of the eye (Fig. 2D, E). The ommatidial hair has a diameter of ˜1.0703 ± 0.3 μm at its base and its length of ˜10.5536 ± 1 μm. At higher magnification, the cornea appears to be smooth and without any corneal nipple (Fig. 2G). The eye makes an angle of 40.853° with the body.

**Figure 2.**
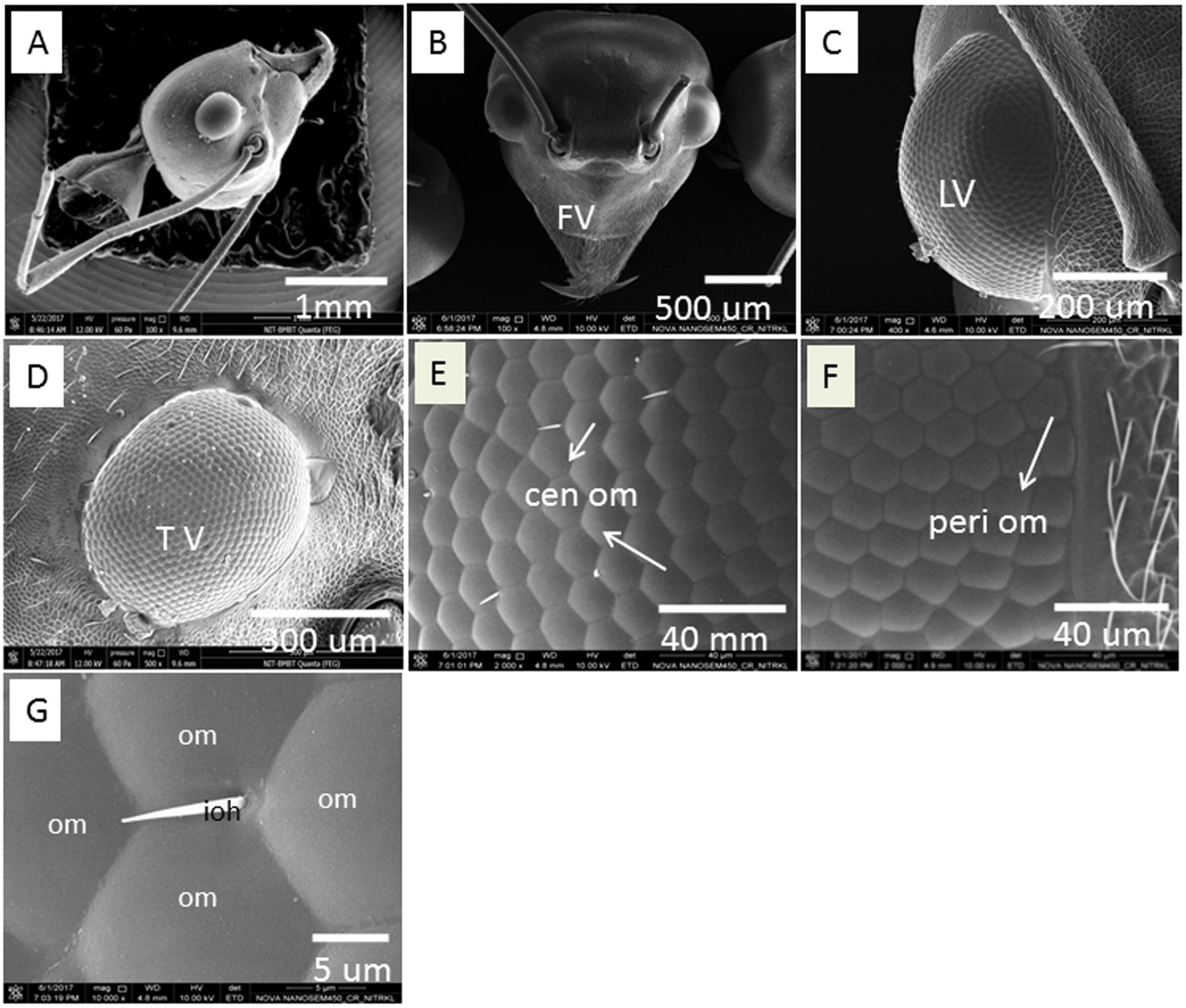
Overview of compound eye of an ant *O. smaragdina*. (A) lateral view of ants head, (B) frontal image (FV) of head, (C) lateral view (LV) of left eye, (D) transverse view (TV) of eye, (E) image of central ommatidia (can om), (F) a view of peripheral ommatidia (peri om), (G) an ommatrichial surrounded by ommatidia.

The current study has chosen the thorax length to measure the size of the ant. The size of the thorax varies from 3.5 to 8.5 μm. The number of facets and surface area of the eye increases progressively as the thorax length increases (Fig. 3A, B). The facet diameter shows a similar trend with an increase of thorax length (Fig. 4A, B). The mean ommatidial diameter was found to be approx. 18.045±2 μm. Furthermore, the facet perimeter and area also increases progressively with thorax length (Fig. 4C, D).

**Figure 3.**
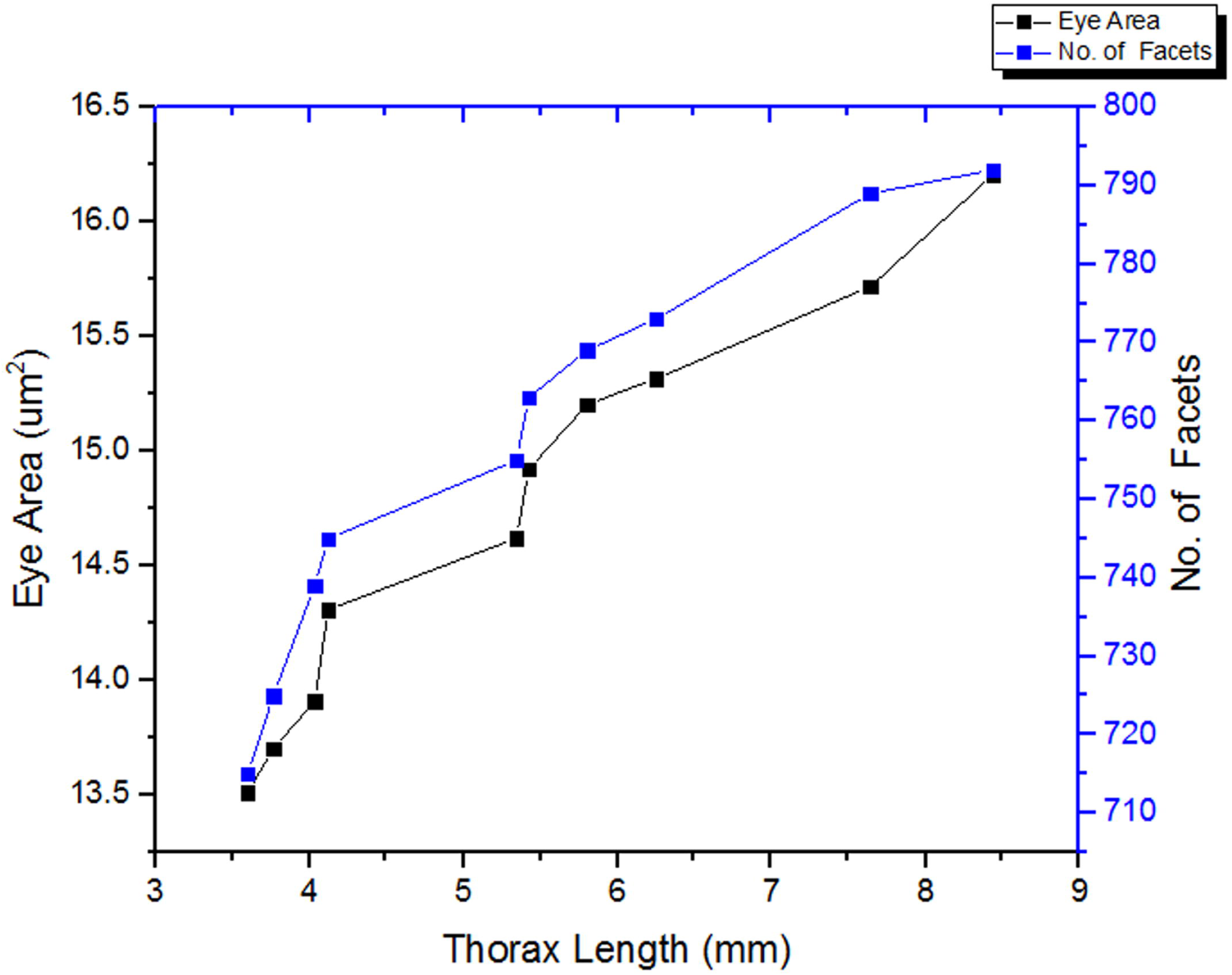
Variation of eye area (μm sq.) and the facet number with the thorax length (mm) Eye area and the number of facets increase significantly with thorax length.

**Figure 4.**
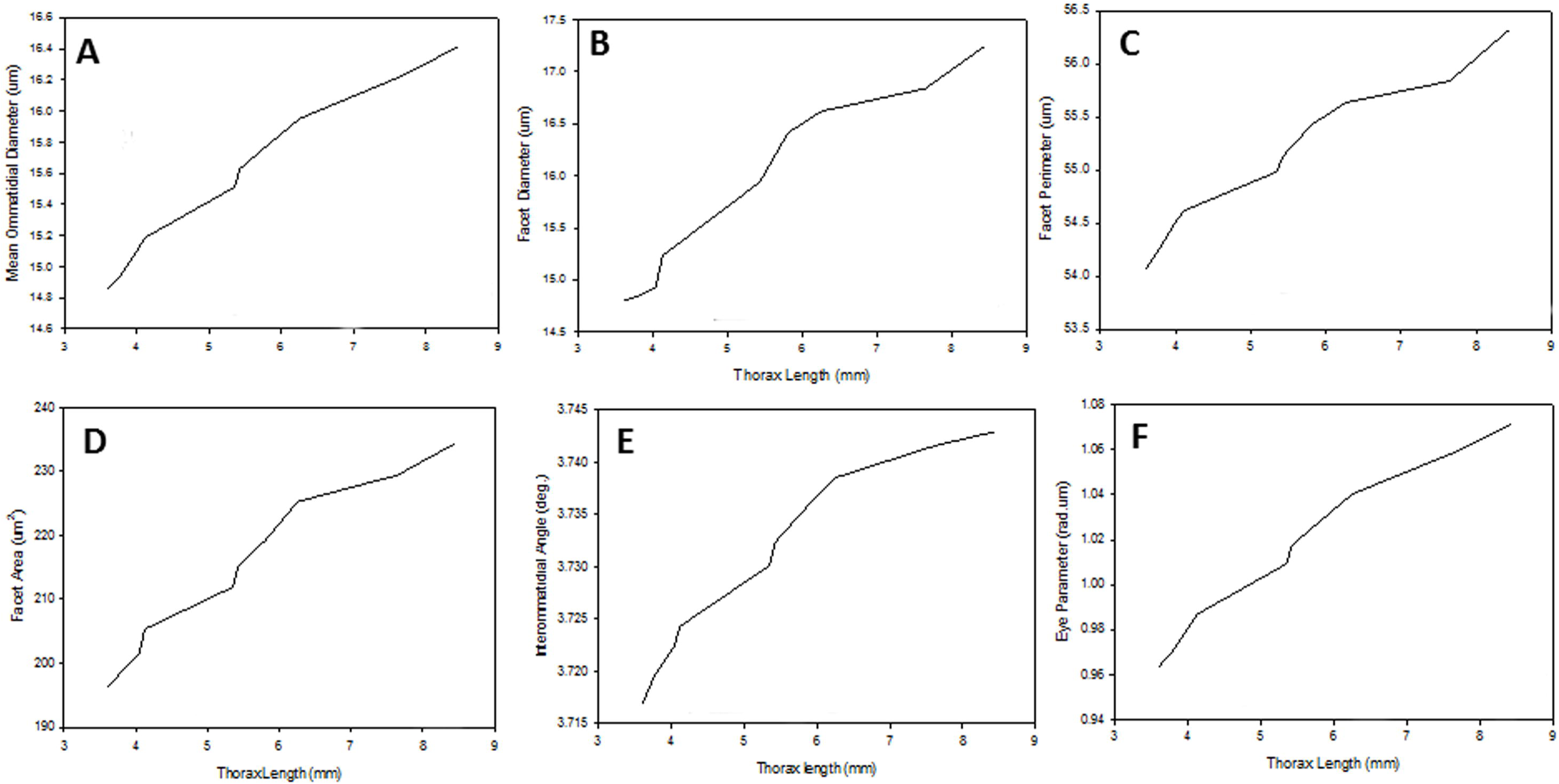
Graphical representations of various parameters of the eye. (A) mean ommatidial diameter (μm), **(B)** facet diameter, **(C)** facet perimeter (μm), **(D)** facet area (μm sq.) with thorax Length (mm) **(E)** interommatidial angle (deg.) **(F)** eye parameter (rad.μm).

To get an idea regarding sensitivity and resolution of the compound eye, the interommatidial angles are calculated. With the increase of thorax length interommatidial angle decreases (Fig. 4E). The interommatidial angle further gives the sampling density of the compound eye. As the thorax length increases, the interommatidial angle increases and hence, higher the potential sampling and resolution of the eye decreases. The value eye parameter (*p*) increases with thorax length which gives the inverse relationship between body length and eye’s resolution.

### Visual discrimination

The visual discrimination of the ants was analyzed in the laboratory using the device mentioned above (Fig. 5A). The ants were introduced into the device through the central part of the hole (Fig. 5 B, C). Ants were allowed to choose various patterns, both small and large field within the device in both the arms of the device (Fig. 5D). First, the ants were allowed to check the light/dark areas. For this experiment, one hand of the arm was kept in light whereas the other was kept in dark. When a light beam was flashed on dark-adapted ants, most of the ants get attracted towards the lighted arm suggesting that the ants prefer light over dark (Fig. 5E). In small filed pattern choice assay more ants preferred upright triangle (Fig. 5F). In the large field pattern choice assay ants are attracted more towards horizontal pattern (Fig. 5G). This pattern recognition experiment suggests that the ants have the ability to recognize various patterns in nature.

**Figure 5.**
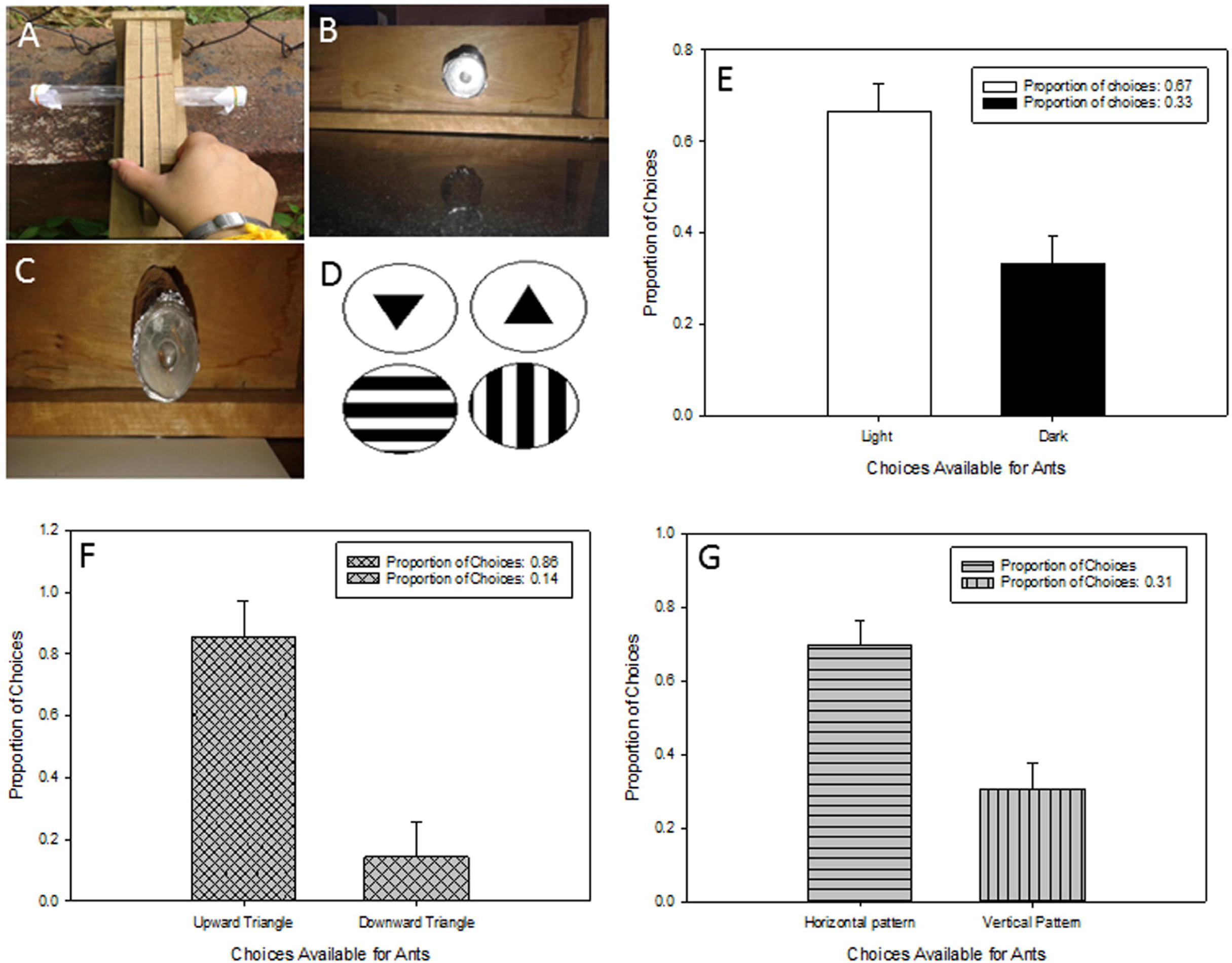
Visual discrimination assay. **(A)** The device used for two-choice assay **(B)** and **(C)** both the arms of the device **(D)** patterns used for visual-discrimination analysis of ants **(E**) ants attracted toward light or dark **(F)** preference of ants towards various triangle **(G)** choice for horizontal and vertical pattern.

## Discussion

The current study correlates the eye structure with its activity time and rhythm of the weaver ant *O. smaragdina*.

### Activity time

Ants set their activity time to reduce the interspecific competitions (Albrecht and Gotelli, 2001; Savolainen and Vepsäläinen, 1988; Whitford and Ettershank, 1975). The activity time is set by choosing the foraging period, microhabitat and type of food taken. Predators are more likely fed on dissimilar prey foraging at a diverse time than herbivore (Gaume and McKey, 1999; Mooney and Tillberg, 2005). Poikilotherms are more sensitive towards temperature, thus the foraging time changes with temperature which makes many ants seasonal (Lynch et al., 1980). Bernstein correlates the foraging activity with temperature which is further linked with periods of more seed production (Bernstein, 1975; Bernstein, 1979). Whiteford found a difference in the foraging activity of two desert ants *Pheidole* and *Pogonomynmex* (Whitford et al., 1999). *Prenolepis*, a violent ant forage during the daytime when the temperature is less in the year and foraging ceases during midsummer (Lynch et al., 1980; Talbot, 1943). Temperature is a key regulator in determining the foraging activity of ants. *Prenolepis* undergo desiccation at 37°C. These physiological restrictions allow them to act at a particular time of the day. Thus *Prenolepis* and *Formica* work at two different times of the day during the different season (Fellers, 1989). Besides food, temperature, and competitions the internal arrangement of the eye provoke them to work in a particular time of the day. For example, the internal arrangement of the day active ant possesses different arrangement than the night active ant (Moser et al., 2004) The structural arrangement further prevents them to be active only at a certain time of the day. The currently studied ant *O. smaragdina* is found to be active only when there is full illumination suggesting the internal arrangement probably allow the animal to be active during the daytime.

### Activity rhythm

Vision has a key role among ants to find food resources in the nearby nest area. Depending on their functionality the ants bear facet numbers from 50 to 4100 in various species (Narendra et al., 2010; Schwarz et al., 2011). Diurnal species like *Camponotous consobrinus* possess 798 facets *(Bernstein and Finn, 1971; Ramirez-Esquivel et al., 2017)*. The current study also reveals the facet number from 710-790. The ants prefer to forage in a group in an environment where the undergrowth is comparatively bare with few proximate landmarks. The number of facets affects the pixel and thus the eye sensitivity. High sensitivity within the eye is not required for navigation (Milford, 2013; Wystrach and Graham, 2012) and target detection. However, the resolution of the eye affects the detection of a small object and detection distance. The compound eye property such as visual fields and the interommatidial angle was calculated from SEM preparation. The number of ommatidia and eye curvature vary in ants and that bring the variations in resolution and visual field (Bernstein and Finn, 1971; Zhang et al., 2007). In the current study also the interommatidial angle and the eye parameter decreases with a body size of the animal. In diurnal ants, the resolution of the eye is very high and sensitivity is less. The eye achieves it by having more number of ommatidia with smaller facet size. Studying foraging behavior and visual tasks in a natural setting give us the information required for the functioning of compound eyes of ants.

#### Facet size and number

The lens diameter of ants varies within the individuals. The lens diameter of the currently studied ant varies from 13 to 18μm. The lens diameter proposes to place *O. smaragdina* with other day active ants like *T. regatulus* (14.7 to 19 μm) and *M. bagoti* 19 μm, *M. croslandi* 18 μm (Greiner et al., 2007; Schwarz et al., 2011). The facet diameter further varies with thorax length. Furthermore, variations were observed in the central and the marginal ommatidia. The marginal ommatidia are irregular in shape. Irregular ommatidia in the marginal region, especially in the dorsal rim area, were observed in several ants such *Ctaglyphis fortis*, *C. consobrinus*, *M. Pyriformis*, *Nothomyrmecia marcopos* and *P. Sokolova* (Narendra et al., 2016; Zeil et al., 2014). The variation observed in the eye shape and size is to achieve various physiological activities for the body. These physiological activities include a selection of mate (Collett and Land, 1975; Snyder, 1977) and detection of prey (Labhart and Nilsson, 1995).

Facet number of the body also increases with the size of the body. Ants like *Cataglyphis albicans* (Zollikofer et al., 1995)*, C. fortis* (Zollikofer et al., 1995), *Camponotus pennsylvanicus* and *Melophorus bagoti* increases facet number with body size to get better resolution (Menke et al., 2009; Menzel, 1972; Perl and Niven, 2016). In species like *Bombus terrestris* and *Solenopsis* sp., and *Formica integroides* both the facet number as well as size increases with body length (Perl and Niven, 2016). As a result compound eye length and area also increases with the size of the body to cope up with the resolution of the eye. Ant-like *Formica rufa* makes its nest with the use of visual clue for its resource allocation(Rosengren, 1977; Skinner, 1980).

Ant worker’s eye size changes with its body size although a negative allometry is established with the eye and body size. In case of wood ants, the eye is further related to the size of the nest (Perl and Niven, 2016). Allometry of the eye size is checked with body size by measuring the facet number, diameter, and size of the eye. In the current study, we also observe the eye size is increasing with body size. Both facet number and size increases with the body size.

#### Visual discrimination

In the current study, the ants prefer upright triangle over the inverted triangle and horizontal lines over the vertical lines. This suggests that the ant has the ability to discriminate between the black and white bands. This further advocate that the ant has the ability to discriminate pattern. The pattern discrimination ability varies from species to species. Although some ants’ species have better recognition ability the others do not possess such ability. In a natural habitat, ants use this property for foraging food or while transferring the larvae from one place to another.

## Conclusions

The current study for the first time reports the activity rhythm and its correlation with the eye structure of weaver ant *O. smaragdina*. The activity rhythm is correlated with the illumination of the surrounding. The activity of the animal is further linked with the structure of the eye. The eye structure share similarity with other diurnal ants. Furthermore, the eye parameter and interommatidial angle also suggest the animal is adopted for the lighted environment. The animal possesses pattern discrimination ability which is required for foraging behavior of the animal.

## Conflict of Interest

We do not have any conflict of interest for this study.

## Ethical approval

This article does not contain any studies with human participants performed by any of the authors.

